# MdBRC1 and MdFT2 Interaction Fine-Tunes Bud Break Regulation in Apple

**DOI:** 10.1101/2025.06.20.660803

**Authors:** Helena Augusto Gioppato, Joan Estevan, Mohamad Al Bolbol, Alexandre Soriano, Julio Garighan, Kwanho Jeong, Céline Georget, Daniela Gómez-Soto, Samer El Khoury, Vítor da Silveira Falavigna, Simon George, Mariano Perales, Fernando Andrés

## Abstract

Winter bud dormancy is a critical adaptive process in temperate fruit trees, safeguarding meristems from freezing temperatures and aligning growth with seasonal cues. Dormancy encompasses two primary phases: endodormancy, where internal signals, particularly elevated abscisic acid (ABA), block growth and necessitate chilling for release; and ecodormancy, where buds regain growth competence but remain quiescent until external conditions are favourable. In apple (*Malus domestica*), we demonstrated that *BRANCHED1* (*MdBRC1*) serves as a central inhibitor of bud growth specifically during ecodormancy. In many plant species, BRC1-like transcription factors integrate environmental and hormonal signals, activating gene networks linked to growth repression, notably those involved in ABA biosynthesis and signalling. Gain-of-function studies in poplar confirmed that *MdBRC1* robustly suppresses shoot growth in trees. Our comparative transcriptomic analyses revealed that MdBRC1 directly regulates a suite of dormancy-associated genes, reinforcing its role as a molecular brake on bud break. Importantly, we show that the apple gene *FLOWERING LOCUS T2* (*MdFT2*) is transcriptionally upregulated after dormancy and might act as a key inducer of bud break. Our data reveal that MdFT2 physically interacts with MdBRC1, reducing MdBRC1 activity during ecodormancy. This antagonistic interaction acts as a molecular switch, facilitating the transition from ecodormancy to active bud growth as spring approaches. Together, these findings uncover a regulatory module that finely tunes bud break timing in apple trees and provide a foundation for breeding strategies to enhance fruit tree resilience and adaptability in the context of climate change.

## RESULTS AND DISCUSSION

### MdBRC1 functions as an inhibitor of bud break and growth

In apple trees (*Malus domestica*), the transition between endo-to eco-dormancy is regulated by a complex gene-regulatory network highly influenced by DORMANCY ASSOCIATED MADS-box (DAMs) and SHORT VEGETATIVE PHASE (SVP) transcription factors ^1^. This regulation culminates in the transcriptional activation of a potential orthologue of the *Arabidopsis thaliana BRANCHED1* (*BRC1*) (MD06G1211100; *MdBRC1*) gene during the ecodormancy ^2^. *BRC1* encodes a class II TEOSINTE BRANCHED1, CYCLOIDEA, PCF (TCP) transcription factor known to shape plant architecture by negatively regulating shoot branching across numerous species ^3–5^. In response to short days, *BRC1*-like genes expressed in the shoot apex of hybrid aspen promote apical growth arrest ^6^. Here, we studied the role of the apple *MdBRC1* during ecodormancy.

The apple genome encodes two closely related putative orthologues of the Arabidopsis *BRC1* gene: *MdTCP18a* (MD06G1211100) and *MdTCP18b* (MD14G1221800) (Supplementary Figure 1). A multiple sequence alignment and phylogenetic analysis of 121 TCP protein sequences, including 24 from *Arabidopsis thaliana* ^7^, 21 from *Arabis alpina* ^8^, 36 from *Populus trichocarpa* ^9^, and 40 from *Malus domestica* ^10,11^, revealed that BRC1 sequences of all four species are grouped closely together within the CYC/TB1 clade, indicating a strong evolutionary relationship (Supplementary Figure 1). Moreover, the high bootstrap values suggest that these genes are highly conserved. The phylogenetic tree suggests that the two closely related BRC1 paralogs in apple, MdTCP18a and MdTCP18b, originated from the genome-wide duplication event that occurred within the apple lineage after its divergence from the common ancestor with *A. alpina* and *A. thaliana* ^11,12^ (Supplementary Figure 1). A sequence similarity analysis between the two apple TCP18 paralogs and the AtBRC1 sequences showed that MdTCP18a shares greater sequence similarity with AtBRC1, suggesting potential conservation of function (Supplementary Figure 2). Based on these findings, MdTCP18a is the most likely BRC1 ortholog in apple.

To get insight into *MdTCP18a* during phenology (hereafter named *MdBRC1*), we investigated its gain-of-function phenotype in a heterologous system using poplar plants (Supplementary Figure 3). Transgenic poplar *MdBRC1* ectopic expressing lines (*MdBRC1#8 and MdBRC1#9*) exhibited a significant delay in bud break compared to wild-type controls (Figures 1A and 1B). This phenotype demonstrates that *MdBRC1* acts as a bud break inhibitor in trees, as *TCP18*/*BRC1* does in hybrid aspen, suggesting the conservation of BRC1 inhibitory role across species ^13^. These findings support the hypothesis that the regulatory mechanisms controlling bud break are functionally conserved, further underscoring the significance of *MdBRC1* in dormancy progression and bud break.

**Figure 1.**
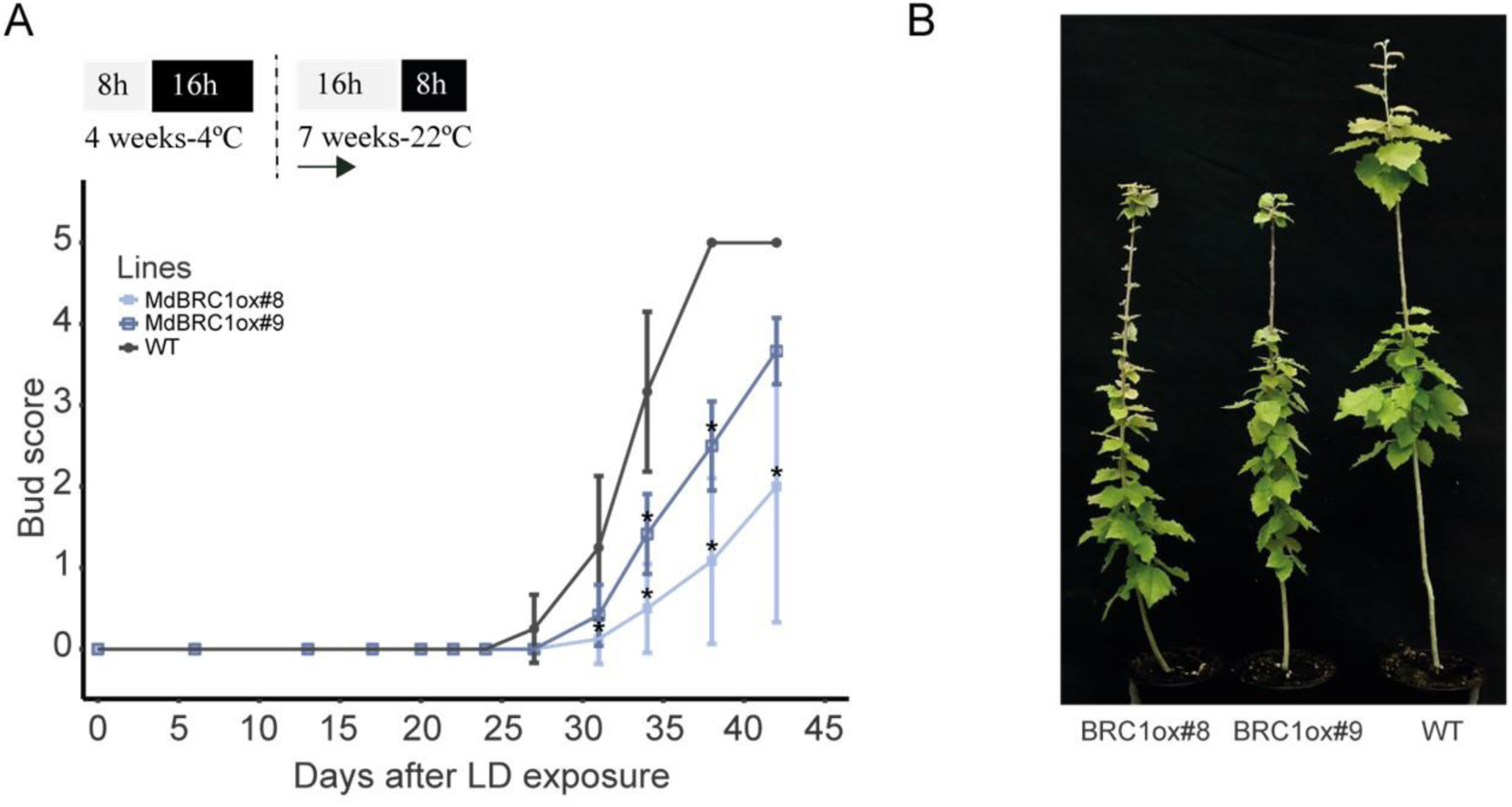
Transgenic poplar plants overexpressing *MdBRC1*. **(A)** Values represent the mean bud score of n=6 plants exposed to LD conditions for 45 days, immediately following 4 weeks of SD and 4°C. Bud break progression was graded on a scale from stage 0 (dormant apical bud) to stage 5 (fully growing apex and elongated internode). Significant differences among lines and the WT were analysed using one-way ANOVA followed by a Tukey test, *p < 0.05. Top panels represent photoperiod (grey boxes mean hours of light; black boxes mean hours of dark) temperature and duration under these conditions. Arrows indicate the start of the experiment; **(B)** Representative photos of the lines taken at the end of the experiment.

### MdBRC1 might play a role in ecodormancy

A previous study reported that *MdBRC1* is transcriptionally activated at the end of endodormancy and remains highly expressed during the ecodormancy phase ^2^. To better understand the function of *MdBRC1* during the transition from endodormancy to ecodormancy and then, during ecodormancy, we analysed global transcriptomic changes in apple buds collected at the early and late stages of both dormancy phases.

The RNA-seq studies revealed global changes in gene expression between the beginning of the endodormancy (time point “C”), the end of the endodormancy (time point “O”), the beginning of the ecodormancy (time point “L”), and the end of the ecodormancy (time point “D”) (Figure 2A, Supplementary Table 1). The pairwise comparison CxD showed that approximately 68% of genes were upregulated at the end of ecodormancy, indicating an important transcriptional activity associated with the transition to growth resumption (Figure 2B, Supplementary Table 2). This finding is consistent with previous reports that explored the transcriptome of apple cultivars during endodormancy and ecodormancy ^14^. Moreover, from the 14,061 differentially expressed genes (DEGs) identified in the time course analysis (Supplementary Table 1), 9,173 genes were included in one of the five clusters (C0-C4) defined based on expression profiles (Supplementary Table 3).

**Figure 2.**
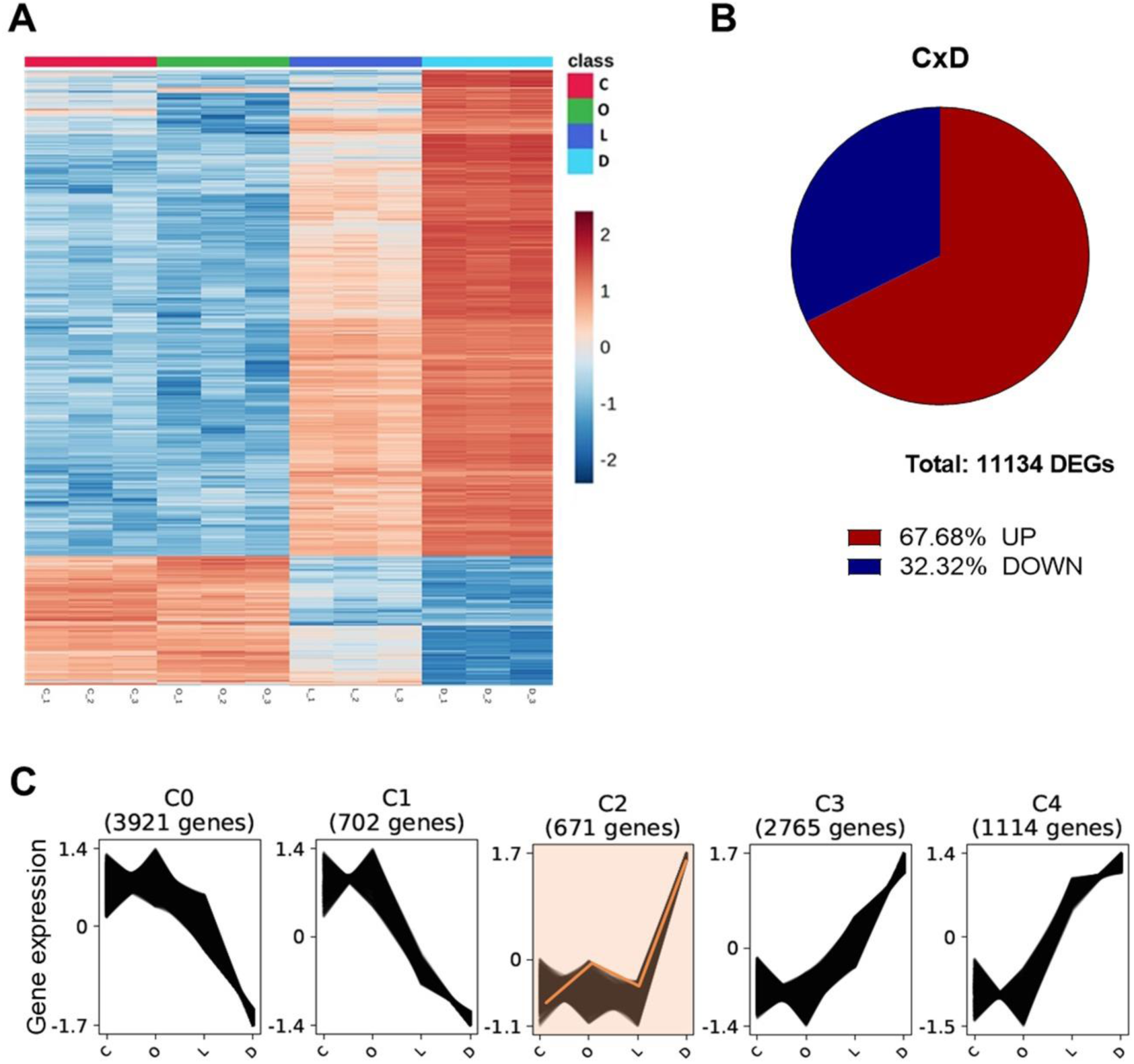
Transcriptomic dynamics during dormancy transitions in apple buds. **(A)** Heatmap showing global transcriptomic changes across four developmental stages: beginning of endodormancy (C), end of endodormancy (O), beginning of ecodormancy (L), and end of ecodormancy (D). Data were log-transformed and normalised using MetaboAnalyst. Hierarchical clustering was performed using Euclidean distance and Ward’s linkage on row-scaled data. Rows represent individual transcripts, and columns represent experimental samples grouped by condition. The colour gradient reflects relative abundance (read counts) levels, with scaling applied by row; **(B)** Pie chart with the percentages of up- and downregulated genes in the pairwise comparison between the beginning of endodormancy (C) and the end of ecodormancy (D) timepoints. Values represent differentially expressed genes (DEGs) which are upregulated or downregulated at the end of ecodormancy (D) when compared to the beginning of endodormancy (C); **(C)** Clustering analysis of differentially expressed genes (DEGs) based on their normalised expression values using Clust. Clusters C0 and C1 include genes downregulated from endo- to ecodormancy, while clusters C2–C4 comprise genes with increasing expression along the same trajectory. The total number of genes per cluster is indicated in parentheses. The orange line represents the expression profile of MdBRC1.

C0 and C1 contained genes being downregulated from endo- to ecodormancy, while clusters C2, C3 and C4 include genes showing the upregulation during the same time course (Figure 2C, Supplementary Table 3). The clusters C0 and C1 presented a downregulation pattern, including genes enriched for Gene Ontology (GO) terms associated with circadian rhythm, photoperiodism, cold and abscisic acid (ABA) responses, ethylene signalling, and ROS metabolism, all considered hallmarks of dormancy-related processes ^15–17^ (Supplementary Table 3). In contrast, clusters C2, C3, and C4 showed upregulation patterns enriched for growth-related terms such as meristem growth, flower development, carbohydrate response, and histone H3-K9 methylation (Figure 2C, Supplementary Table 3) ^18,19^.

*MdBRC1* was found to be upregulated during apple bud ecodormancy in the RNA-seq analysis. It was included in cluster C2, which showed significant enrichment for the terms carbohydrate catabolic process (GO:0016052), small molecule metabolic process (GO:0044281), and response to cadmium ion (GO:0046686) (Supplementary Table 3). The induction of genes involved in carbohydrate and small molecule metabolism suggests increased energy demand and resource mobilisation to support cellular activity under changing physiological conditions ^20,21^. The enrichment of cadmium response genes, despite the absence of cadmium, may indicate the activation of general stress pathways ^22^. The expression pattern of *MdBRC1* points to a role in regulating ecodormancy by affecting the expression of genes across different clusters.

### MdBRC1 affects the ABA signalling pathway and its activity is antagonised at ecodormancy

To better understand the MdBRC1 molecular function in controlling ecodormancy and bud break, we next identified its transcriptional targets and their regulation using a glucocorticoid receptor (GR) system coupled with RNA-seq. To this end, we produced apple calli transformed with a construct overexpressing *MdBRC1-GR* from a *35S* promoter. Using this system, we identified a total of 277 DEGs (adj. p≤ 0.05) regulated by MdBRC1 activity. Of these, 206 (74.4%) were upregulated and 71 (25.6%) downregulated upon dexamethasone (DEX) treatment (Figures 3A and 3B). GO analysis revealed that the functions of MdBRC1 target genes were enriched in processes such as photosynthesis, transcription, proton transport, ABA response, and ROS metabolism (Figure 3C, Supplementary Tables 4 and 5), which is consistent with those described for the BRC1 ^23^.

**Figure 3.**
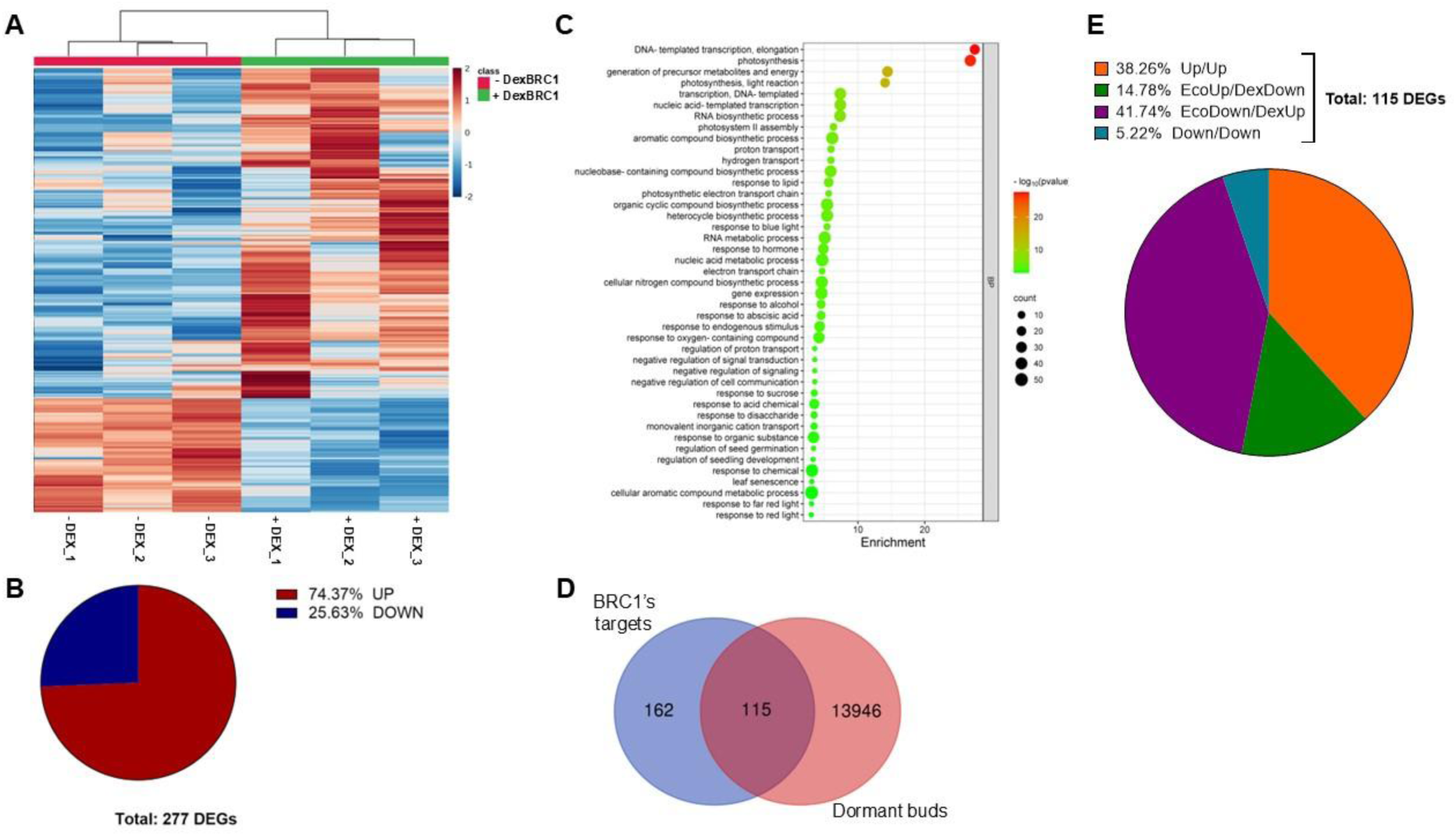
MdBRC1 transcriptional targets and their relation to bud dormancy. **(A)** Heatmap showing differentially expressed genes (DEGs) in apple calli expressing MdBRC1-GR treated with dexamethasone (DEX), generated using MetaboAnalyst. Gene expression data were log-transformed and normalised before hierarchical clustering (Euclidean distance, Ward’s method); **(B)** Pie chart showing the percentages of MdBRC1 transcriptional targets: 74.4% were upregulated, corresponding to 206 genes (red) and 25.6% were downregulated, corresponding to 71 genes (blue); **(C)** Gene Ontology (GO) enrichment analysis of MdBRC1 target genes was performed using agriGO, applying a significance threshold of FDR ≤ 0.05. The analysis revealed significantly over-represented biological processes, including photosynthesis (GO:0015979), response to abscisic acid (GO:0009737), and response to oxygen-containing compound (GO:1901700); **(D)** Venn diagram showing the overlap between MdBRC1 transcriptional targets (blue) and dormancy-related DEGs (red), identifying 115 Dormancy-Related BRC1 Targets (DRBTs; the intersection); **(E)** Pie chart illustrating the regulation of DRBT genes at the end of ecodormancy and the BRC1 target genes datasets. Orange and blue portions represent the group of genes that have the same regulation in both conditions (up/up and down/down, respectively), while green and purple portions represent genes with contrasting regulation profiles (EcoUp/DexDown and EdoDown/DexUp, respectively).

To identify targets of MdBRC1 involved in dormancy regulation, we compared the MdBRC1 targets with the DEGs found in the transcriptomics of dormant buds. The common genes for the two lists were considered Dormancy-Related BRC1 Targets (DRBTs). We identified 115 DRBTs, out of which 92 were upregulated and 23 downregulated by *MdBRC1* (Figures 3C and 3D). Remarkably, MdBRC1 induceed the expression of several ABA biosynthetic genes, such as *nine-cis-epoxycarotenoid dioxygenase 3* (*NCED3*) and *NCED5* (Supplementary Table 6).

*NCED3* and *NCED5* encode 9-cis-epoxycarotenoid dioxygenases, catalysing the rate-limiting oxidative cleavage of 9-cis-epoxycarotenoids to xanthoxin in the ABA biosynthetic pathway ^1,24,25^. Similarly, *NCDE*-encoding genes are direct targets of the BRC1 transcription factor in Arabidopsis ^23^. Our results indicate that MdBRC1 not only promotes ABA biosynthesis by inducing the expression of *NCED3* and *NCED5* but also modulates ABA sensitivity through the transcriptional activation of *HAI3* (*highly ABA-induced PP2C gene 3*) genes, which encode a clade A PP2C phosphatase that negatively regulates ABA signalling ^26,27^. This suggests that MdBRC1 coordinates both the production and fine-tuning of ABA responses. We observed a similar regulation in poplar plants overexpressing *MdBRC1* (Supplementary Figure 4).

Unexpectedly, several DRBT genes that were either upregulated or downregulated by MdBRC1 activity in the DEX experiment exhibited the opposite expression pattern during ecodormancy, a stage in which *MdBRC1* is highly expressed (Figure 3E, Supplementary Table 6). This suggests that, although MdBRC1 regulates the transcription of these DRBT genes in calli, its ability to modulate their expression during ecodormancy may be influenced by additional factor(s). Notably, many of these genes encoded ABA biosynthetic enzymes and transcriptional regulators, which were induced by MdBRC1 in the DEX experiment and downregulated at the end of ecodormancy (e.g. *MdNCED3, MdNCED5, MdHAI3a/b/c and MdABI 1/5*, Supplementary Table 6).

### MdFT2 antagonises MdBRC1 activity

Previous studies demonstrated that the transition from growth cessation to dormancy, and subsequently to growth resumption, is regulated by the floral integrator gene *FLOWERING LOCUS T* (*FT*) in *Populus* species ^28–30^. In Arabidopsis, FT protein has been shown to physically interact with BRC1, potentially modulating its activity ^31^. Poplar has two *FT* paralogs (*FT-like* genes) with distinct developmental roles: *FT1* determines flowering induction and dormancy release in response to long days, and *FT2* promotes shoot apex development under long days in the growing season and triggers growth cessation in response to short days in autumn ^29,32,33^. BRC1 physically interacts with FT2 and inhibits its activity, leading to growth cessation ^34^.

In the apple genome, there are also two *FT* paralogs, *MdFT1* and *MdFT2*, whose seasonal expression patterns suggest roles in floral transition during summer and bud break in spring, respectively ^35,36^. These functions are further supported by transgenic studies showing that overexpression of these genes induces early flowering and overrides bud dormancy in apple trees ^37^. Therefore, the apple FT-like proteins are strong candidates for modulating MdBRC1 activity during ecodormancy.

In our RNA-seq analysis of dormant buds, we observed changes in the expression of both *MdFT-like* genes, although these differences were not statistically significant with the criteria we imposed (adj. p≤ 0.05). *MdFT1* showed peak expression at the onset of ecodormancy, with a gradual decline toward the end of this phase. In contrast, *MdFT2* expression peaked at the end of ecodormancy, coinciding with the highest levels of *MdBRC1* expression (Figure 4A). We could confirm this pattern of expression using RT-qPCR on a time-course experiment during the dormancy cycle (Figure 4B). These results suggest that *MdFT2* is the more likely *MdFT-like* candidate to influence MdBRC1 function during ecodormancy. Furthermore, our protein–protein interaction assays revealed that MdBRC1 physically interacts with both apple FT paralogues, suggesting that the molecular interaction underlying BRC1–FT regulatory relationships, previously described in Arabidopsis and poplar, are conserved in fruit tree species as well (Figure 4C).

**Figure 4.**
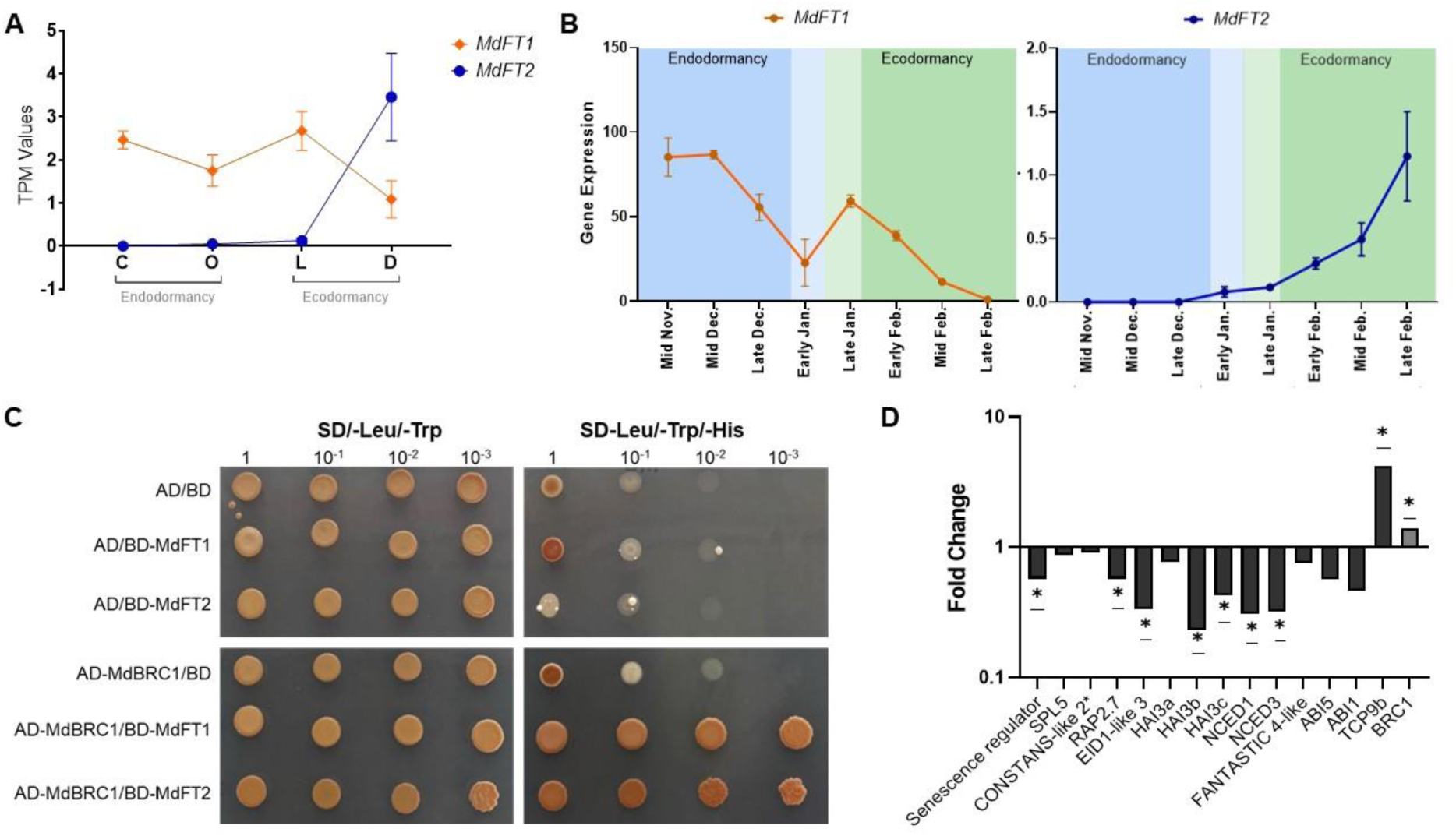
Expression dynamics and protein interactions between MdFT-like and MdBRC1 during bud dormancy. **(A)** Relative expression levels of MdFT1 and MdFT2 across dormancy at the beginning and end of endodormancy (C and O, respectively), measured by RNA-seq. Data are presented as mean TPM (Transcripts Per Million) ± SEM (n = 3 biological replicates). Although these trends were consistent, they did not meet statistical significance under the adjusted P ≤ 0.05 threshold; **(B)** Relative expression of MdFT1 and MdFT2 measured by RT-qPCR over time. Expression levels are shown as means ± SEM (n = 6). Samples were collected from November 2016 to February 2017; **(C)** Yeast two-hybrid (Y2H) assay showing that MdBRC1 physically interacts with both MdFT-like proteins. MdFT1 and MdFT2 in the pDEST32 were used as baits, and MdBRC1 in the pDEST22 was used as prey. Co-transformation of pDEST32 and pDEST22 empty vectors and co-transformations of the empty vector with a pDEST vector containing MdFT1, MdFT2 or MdBRC1 were used as negative controls. Yeast cells grown on an SD plate deficient in Tryptophan and Leucine (SD/-Trp/-Leu) indicate correct co-transformation. Yeast cells grown on an SD plate deficient in Tryptophan, Leucine, and Histidine (SD/-Trp/-Leu/-His), supplemented with 5 mM 3-Amino-1,2,4-triazole (3AT), indicate the interactions between MdBRC1 and MdFT1 and MdFT2; **(D)** Fold change in the expression of MdBRC1 target genes in apple seedlings overexpressing *35S::MdBRC1* alone or in combination with *35S::MdFT2*, relative to seedlings transformed with the empty p19 vector (control). Due to differences in MdBRC1 expression between the two conditions, target gene expression levels were normalised to the relative expression of MdBRC1 in each treatment, enabling the comparison of MdBRC1 regulatory activity on its targets and the evaluation of whether MdFT2 modulates this effect. Plotted values are fold-change means (n = 6 biological replicates). Bars marked with an asterisk indicate statistically significant differences (*t*-test, *p* ≤ 0.05).

To further understand how the interaction between MdFT2 and MdBRC1 could affect the function of the latter, we performed transient overexpression experiments in apple seedlings. In these experiments, we transformed apple plantlets with constructs that overexpressed *MdBRC1* and *MdFT2* individually and simultaneously. Then, we evaluated the expression levels of several DRBT genes through RT-qPCR, in order to test the hypothesis that MdFT2 disturbs MdBRC1 function. Our results confirmed that, in most cases, such as for *NCDE* and *HAI3* genes, the ability of MdBRC1 to regulate the transcriptional levels of DRBT genes was impaired in the presence of high levels of *MdFT2* (Figure 4E).

Altogether, these findings strongly suggest that the interaction between MdFT2 and MdBRC1 negatively affects MdBRC1’s transcriptional activity. This finding complements previous reports that have highlighted the role of BRC1–FT interactions in regulating growth cessation in trees ^6^. In addition to FT, Arabidopsis class I transcription factors TCP14 and TCP15 antagonize BRC1 function. Specifically, TCP15 physically interacts with BRC1, sequestering it from chromatin and thereby restricting its capacity to activate transcription of downstream target genes ^38^. In our RNA-seq experiment, only *MdTCP15b* was differentially expressed in dormant buds (Supplementary Figure 5). However, since *MdTCP15b* is downregulated during ecodormancy, it is unlikely that the MdTCP15b protein interferes with MdBRC1 function in the manner we propose for MdFT2.

Thus, based on our results, we propose a model of finely tuned bud break control in apple trees, in which MdBRC1 contributes to maintaining buds in an ecodormant state until environmental conditions become favourable for growth. In this model, MdFT2 may function as an integrator of spring-associated environmental cues, such as rising temperatures, to counteract MdBRC1’s inhibitory effect on bud break. The interaction between MdBRC1 and MdFT2 establishes a regulatory balance between ecodormancy and bud activation. Under non-permissive conditions (e.g., low temperatures), elevated *MdBRC1* levels help sustain ecodormancy by inducing ABA biosynthesis and regulating its signalling pathways. As conditions shift to more favourable ones (e.g., warmer temperatures), *MdFT2* expression increases, and the MdFT2 protein interacts with MdBRC1. Once MdFT2 accumulates beyond a certain threshold, this interaction inhibits MdBRC1’s ability to regulate its downstream targets (e.g., *ABI1*, *ABI5*, etc.), thereby promoting bud break (Figure 5).

**Figure 5.**
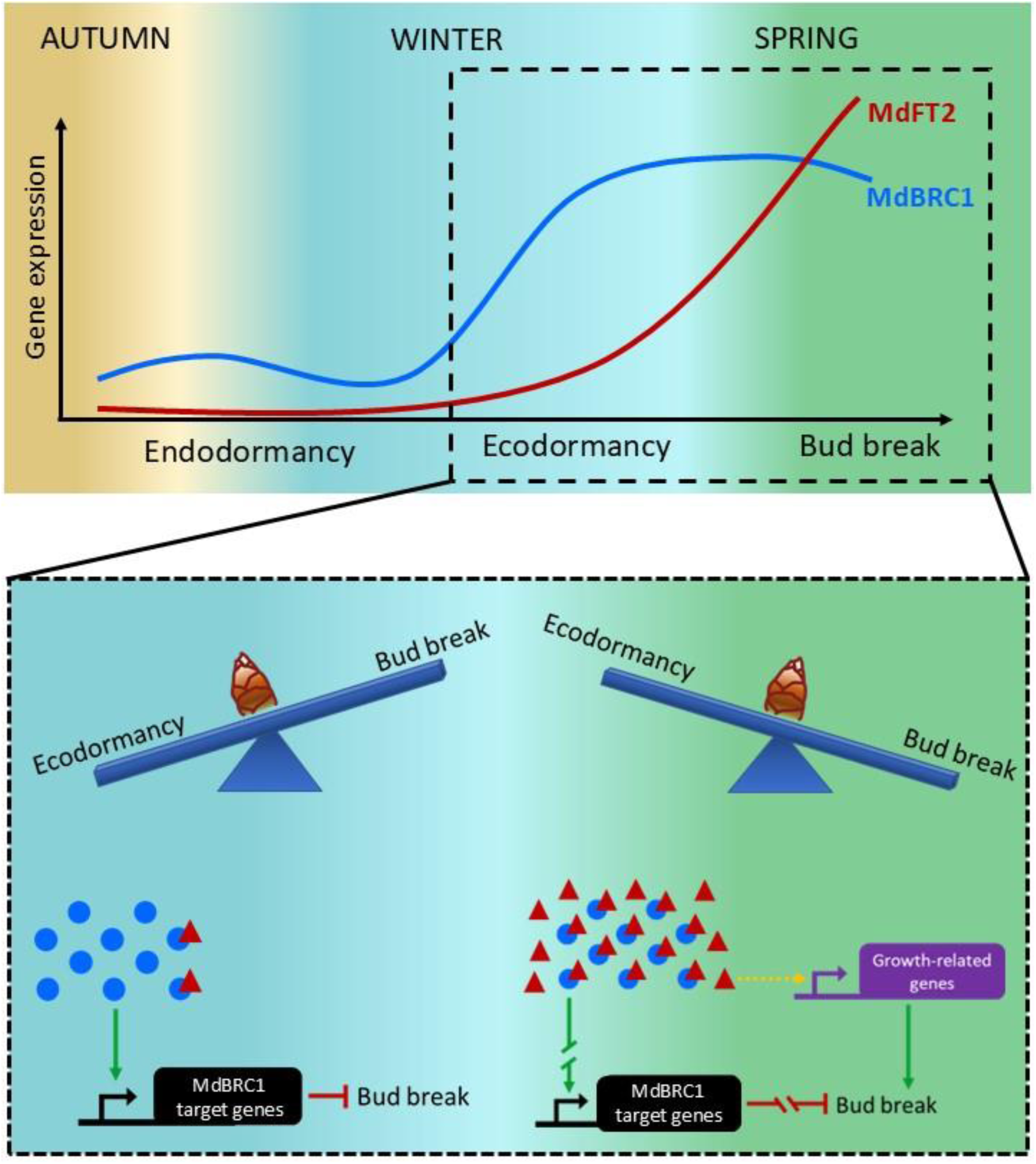
Proposed model for the regulatory interplay between MdBRC1 and MdFT2 in the control of bud dormancy and activation. Under non-inductive conditions (e.g., low temperature), high *MdBRC1* (blue line) transcriptional levels promote ecodormancy maintenance by inducing ABA biosynthesis and regulating its signalling components. As spring-associated signals (e.g., high temperatures) increase *MdFT2* expression (red line), its protein (red triangles) interacts with MdBRC1 proteins (blue circles), progressively reducing its activity. Once MdFT2 accumulates beyond a threshold, it suppresses MdBRC1’s regulation of downstream targets (e.g., *NCED3*, *ABI5*, *HAI3*), thus promoting bud break. Green and red arrows indicate induction and inhibition, respectively. The yellow dashed arrow indicates an indirect induction effect.

## CONCLUSION

In conclusion, our study advances the understanding of the complex molecular mechanisms regulating dormancy and bud break in apple trees. We demonstrate that MdBRC1 plays a crucial role in dormancy control, particularly during ecodormancy, by supporting the growth-inhibitory activity of the hormone ABA. It has been shown that ABA biosynthesis is induced by cold and by the activity of DAM transcription factors at the onset of endodormancy ^6,39^. These transcription factors promote *MdBRC1* expression toward the end of endodormancy. In turn, MdBRC1 reinforces ABA activity by regulating genes involved in its biosynthesis and signalling. We propose a mechanism that resolves the apparent paradox of high *MdBRC1* levels and the need for reduced ABA levels to trigger bud break in spring. This model builds on a well-established regulatory module involving the interaction between FT and BRC1. While this module has been shown to control bud break and flowering in herbaceous species ^3,31^, as well as growth cessation in trees ^13,40^, our findings extend this knowledge by demonstrating that FT2 inhibits the ABA-dependent bud break-suppressing activity of MdBRC1 in fruit trees. This insight provides a foundation for developing innovative strategies to enhance the resilience of apple and other temperate fruit tree species in the face of climate change.

## STAR METHODS

- **RESOURCE AVAILABILITY**

`o` **Lead contact**
`o` **Material availability**
`o` **Data and code availability**
- **EXPERIMENTAL MODEL AND SUBJECT DETAIL**
- **METHOD DETAIL**

`o` **Plant material and growth conditions**
`o` **Generation of plasmids for plant transformation and cloning**
`o` **Plant transformation**
`o` **Glucocorticoid Receptor assay**
`o` **RNA-seq library construction, sequencing and analysis**
`o` **Sequence identification and phylogenetic analysis**
`o` **RNA isolation and RT-qPCR analysis**
`o` **Yeast Two-Hybrid (Y2H)**
- **QUANTIFICATION AND STATISTICAL ANALYSIS**

## STAR METHODS RESOURCE AVAILABILITY

### Lead contact

Further information and requests for resources and reagents should be directed to and will be fulfilled by the Lead Contact, Fernando Andrés. This study has not generated any new reagents.

### Material availability

All unique/stable reagents generated in this study are available from the lead contact with a completed Materials Transfer Agreement.

### Data and code availability

- RNA-seq data will be deposited at the NCBI and are publicly available as of the date of publication. Accession numbers will be listed in the key resources table.
- Any additional information required to reanalyse the data reported in this paper is available from the lead contact upon request.

## EXPERIMENTAL MODEL AND SUBJECT DETAIL

*Malus domestica* cvs. “Golden Delicious” and “Gala” were used as the experimental model.

## METHOD DETAIL

### Plant material and growth conditions

#### For dormancy RNA-seq

Buds from the apple cultivar “Golden delicious” collected between 2018 and 2019 were used for the RNA-seq. Three buds from short shoots (spurs of more than two years) per date and tree were collected. A total of nine buds from three trees were pooled together to compose an independent biological sample. In this way, three biological replicates from the same nine trees were collected at each date.

#### For calli RNA-seq

The apple cultivar ‘Gala’ was used in this study. In vitro cuttings of apple were subcultured at one-month intervals on a micropropagation medium, a modified Lepoivre medium ^41^ supplemented with 0.5 mg/L 6-benzylaminopurine and 0.1 mg/L indole-3-butyric acid, at 22°C under a 16-h light (80 μmol m⁻² s⁻¹)/8-h dark photoperiod.

#### Generation of plasmids for plant transformation and cloning

For the overexpression assay, *MdBRC1* and *MdFT2* coding sequences (CDS) were cloned into the pDONR207 vector ^42^, using the Gateway system (Thermo Fisher Scientific). Then, they were recombined into the pK7WG2D binary vector ^43^, which carries the *in planta* selection marker *35S*::*GFP* (*GREEN FLUORESCENT PROTEIN*) and an additional constitutive-expression promoter CaMV *35S* to drive the expression of *MdBRC1* and *MdFT2*, generating the plasmids *35S::MdBRC1* and *35S::MdFT2*.

For the GR inducible system, the *MdBRC1* pDONR207 vector was recombined into a modified pK7WG2D binary vector carrying the GR protein at N-terminus position to generate *35S::MdBRC1*-*GR*. All vectors were confirmed by sequencing and used in the transformation in *A. tumefaciens* strain EHA105 ^44^.

To generate *MdBRC1* overexpressing poplar lines, *MdBRC1* coding sequences (CDS) were cloned into the pDONR207 were recombined into the pMDC83 binary vector ^45^, which carries the *in planta* selection marker *Hygromycin* and an additional constitutive-expression promoter CaMV *35S* to drive the expression of *MdBRC1,* generating the binary plasmid *35S::MdBRC1*.

#### Overexpression of *MdBRC1* and *MdFT2* in apple seedlings

The previously described apple transformation protocol ^46^ was modified as follows: The inoculum was a mix of *Agrobacterium* transformed with *35S::MdBRC1* and/or *35S::MdFT2* and a strain transformed with a gene coding the p19 protein, a suppressor of gene silencing ^47^, at OD_600_ = 5 and 2.5, respectively. The cultures were resuspended in an induction medium containing 150 μM acetosyringone ^48^ and incubated at room temperature with gentle agitation for 3-4 h and then supplemented with 0.002% (v/v) Silwet-77 before use. *In vitro* cuttings were totally submerged in the inoculum and vacuum infiltrated for 1 min at -0.09 mPa. Shoots were then rinsed in three successive baths of sterile water, dried on sterile filter paper and placed in a growth chamber on a micropropagation medium without antibiotics.

Four conditions were used: cuttings transformed with p19, *35S::MdBRC1*, *35S::MdFT2*, and *35S::MdBRC1* and *35S::MdFT2* simultaneously. For each condition, 12 plants were agro-infiltrated. Six seedlings per condition were harvested at 2 and 5 days post-transformation and then immediately frozen in liquid nitrogen and stored at -80°C until analysis.

#### Overexpression of *MdBRC1* in Poplar

The hybrid poplar Populus tremula × alba INRA clone 717 1B4 was used as the experimental model. Poplar plantlets were grown in vitro in Murashige and Skoog medium 1B (pH 5.7) supplemented with 2% sucrose and with indole acetic acid (0.5 mg/L) and indole butyric acid (0,5 mg/L) containing 0.7% (w/v) plant agar under long days (LD) 16h light/ 8h dark and 22 °C conditions. The binary plasmid *35S::MdBRC1*. was transformed into *Agrobacterium tumefaciens* strain GV3101/pMP90 ^49^. Next, poplar explants were transformed via an Agrobacterium-mediated protocol described previously^50^. Selection was conducted in hygromycin-containing medium, and once plantlets were regenerated, the gene expression levels of *MdBRC1* overexpressing lines were analysed by quantitative reverse transcription-polymerase chain reaction (RT-qPCR) in the independent transformed individuals (lines) and in wild-type individuals. The *MdBRC1#8 and MdBRC1#9* lines were selected to conduct phenological experiments.

#### Phenological assays of poplar *MdBRC1* overexpressing lines

For the phenological assays, in vitro cultivated poplars of wild type (WT), *MdBRC1#8 and MdBRC1#9* lines were transferred to blond peat pots (pH 4.5). Plants were grown under LD and 22 °C for 4 weeks. Plants were fertilised once every 2 weeks with a solution of 1 g l−1 Peters Professional 20-20-20 in LD conditions and 20-10-20 in short day (SD, 8h light/16h dark) conditions (Comercial Química Massó, Barcelona, Spain). Growth cessation and bud set were induced by exposing plants to SD conditions (8h light/16h 16h dark) and 22°C for 10 weeks. Bud set progression was graded by scoring from stage 3 (fully growing apex) to stage 0 (fully formed apical bud) according to Rohde et al., 2010. Winter conditions are simulated by treating plants under SD and 4 °C for 6 weeks to satisfy their chilling requirement. In this study, we conducted an insufficient chilling treatment involving SD and 4°C for only 3 weeks to monitor the early release from dormancy in the WT, *MdBRC1#8 and MdBRC1#9* lines. Finally, plants were transferred back to LD and 22 °C to monitor bud break. The bud break was scored according to the six developmental stages (from 0 to 5) according to Johansson et al. (2014) ^51^.

#### Production of transformed calli with *35S::MdBRC1-GR* and chemical induction

Apple leaves were transformed with *Agrobacterium* containing the construct *35S::MdBRC1*-*GR*, according to the protocol described by Estevan, J. et al. (2020) ^52^. Calli derived from apple leaf explants were arranged in two groups, the ‘Control’ (30 non-treated samples) and ‘Treated’ (30 treated samples). Control samples were treated with a mock solvent, while Treated samples were submitted to cycloheximide (CHX) followed by a dexamethasone (DEX) treatment (see Estevan, J. et al., 2020).

### RNA-seq library construction, sequencing and analysis

#### RNA extraction and library construction for RNA-seq analysis

Total RNA was isolated using the Spectrum Plant Total RNA kit (Sigma-Aldrich), the integrity of the total RNA was analysed using a 4200 TapeStation instrument (Agilent Technologies, Santa Clara, California, USA), while the RNA quantification was performed with a NanoQuant instrument (Infinite 200 NanoQuant; Tecan, Mannedorf, Switzerland). The RNA was subjected to a DNase treatment using a Turbo DNA-free kit (Thermo Fisher Scientific). Libraries were constructed with the Illumina’s TruSeq Stranded mRNA kit (Illumina) and sequenced on two lanes of an S4 flow cell in paired-end 150 nt (NovaSeq6000, Illumina) at MGX (Montpellier, France).

#### Read cleaning and mapping

Cutadapt (v4.1) ^53^ was used to find and remove Illumina Trueseq adapter sequences and to remove low-quality base pairs in both 5’ and 3’ of each read based on a quality cutoff of 20 ^45^. All processed reads with final lengths smaller than 50 were discarded. Cleaned reads have been mapped using Hisat2 (v2.2.1) ^54^ onto the GDDH13 V1.1 reference genome (https://iris.angers.inra.fr/gddh13/the-apple-genome-downloads.html) using the provided annotation. The sequence of “MdoChr04g0019" from Malus x domestica ’Hanfu’ was downloaded from https://www.rosaceae.org, extended to 100pb in both 5’ and 3’ and has been added to the reference genome. The Hisat2 options “--sensitive”, “--rna-strandness RF” and “--max-intronlen 25000” have been used ^54^.

#### Time course analysis

For the time course analysis of the dormancy RNA-seq, read quantification was made using Kallisto (v.0.44.0) with 100 bootstraps, returning TPM (Transcripts Per Million) values representing the transcripts’ abundance ^55^. R package sleuth (v. 0.29.0) was used to analyze this dataset for the time course differential expression analysis ^55^. It uses the likelihood ratio test (LRT) and splines regression to select genes that follow a differential pattern through time, excluding the possibility of noise.

#### Pairwise comparisons

For the pairwise comparison of the RNA-seq data from the dexamethasone experiment, read counting has been performed using FeatureCount from the subread package (v2.0.1) using the “-M --fraction” parameter to handle multi-mapping reads ^56^. Next, read counts were processed using the graphical interface DIANE (Dashboard for the Inference and Analysis of Networks from Expression data) ^57^. Data was normalised with the TMM method (Trimmed Mean of M values). After filtering, only genes with a sum of reads equal to or greater than 10 in both conditions were conserved. Different expression analysis used a cutoff of Log2FC = 1 and adj. p ≤ 0.05.

#### Gene Ontology (GO) enrichment analysis

The GO enrichment analysis of the differentially expressed genes (DEGs) was performed in agriGO (v2.0) ^58^.

#### Clust

To seek genes with similar expression patterns along the phase transitions, we have made a co-expression analysis with the software Clust (v.1.10.12) using as input the set of significantly differentially expressed genes in the time course comparison ^59^. Clust received 1 dataset with 14 061 unique genes. After filtering, 14 061 genes made it to the clustering step. Clust generated 5 clusters of genes, which in total include 9 173 genes. The smallest cluster includes 183 genes, the largest cluster includes 3 921 genes, and the average cluster size is 671 genes.

#### Sequence identification and phylogenetic analysis

*Malus domestica* TCP sequences were obtained from the GDR database ^60^ and renamed based on updates by Tabarelli et al. (2022) ^10^. The sequences from *Arabis alpina* and *Populus trichocarpa* were retrieved from PlantTFDB ^61^, with poplar TCP genes following Ma et al. (2016) ^9^. Sequences from Arabidopsis thaliana were sourced from The Arabidopsis Information Resource (TAIR, https://www.arabidopsis.org/). Full-length amino acid sequences of TCP proteins aligned using the L-INS-I algorithm of MAFFT v7 software ^62,63^. The best-fit model, selected according to the Akaike criterion, was JTT+F+G4, which was then used for phylogenetic reconstruction. Phylogenetic reconstruction was performed using the Maximum Likelihood method in IQ-TREE v1.5.4 ^64^, with bootstrap support assessed using 1000 replicates. Additionally, tree branches were tested using the SH-like aLRT method with 1000 replicates.

#### Sample preparation for apple RT-qPCR analysis

The same RNA samples of dormant buds used for RNA-seq were used for gene expression analysis by RT-qPCR. The SuperScript III First-Strand Synthesis System (Thermo Fisher Scientific, Waltham, MA, USA) was used for cDNA synthesis. Total RNA was isolated using the Spectrum Plant Total RNA kit (Sigma-Aldrich), and the SuperScript III First-Strand Synthesis System (Thermo Fisher Scientific, Waltham, MA, USA) was used for cDNA synthesis. Real-time polymerase chain reaction was performed using the LightCycler 480 instrument (Roche, Mannheim, Germany). The primers used are listed in Supplementary Table 7. *MdWD40* was used as a reference gene ^65^.

For gene expression analysis of MdBRC1 target genes in apple seedlings overexpressing *MdBRC1* alone or in combination with *MdFT2*, we first confirmed the transformation efficiency by comparing *MdBRC1* and *MdFT2* expression levels in seedlings transformed with the empty p19 vector (control), *35S::MdBRC1*, *35S::MdFT2* and the combined (*35S::MdBRC1* + *35S::MdFT2*). Next, the expression of known MdBRC1 target genes in both *35S::MdBRC1* and (*35S::MdBRC1* + *35S::MdFT2*) seedlings was calculated relative to the control (p19). However, due to the different MdBRC1 expression levels in these two conditions, the expression of MdBRC1 target genes was normalised to the relative expression of MdBRC1 in each condition. This normalisation allowed the comparison of regulatory effects of MdBRC1 on its targets, independently of variation in its expression, and to assess whether MdFT2 modulates this effect.

#### RNA extraction and RT-qPCR analysis of poplar overexpression lines

For comparative gene expression analysis of MdBRC1#8 and MdBRC1#9 apical buds, samples were collected and immediately frozen. Two biological replicates were collected for this purpose. Total RNA was extracted from the ground buds using a cetyltrimethylammonium bromide (CTAB) extraction buffer supplemented with 2% β-mercaptoethanol. The mixture was incubated at 65∘°C for 5-8 minutes, followed by three washes with chloroform. Next, 7.5 M LiCl2:50mM EDTA was added, and the samples were precipitated overnight at 4°C. The following day, RNA was collected by centrifugation at 10,000 RPM for 20 minutes. The supernatant was discarded, and the pellet was resuspended in NR buffer. RNA was then purified according to the manufacturer’s instructions using the NZY RNA purification kit (NZYTech, Lisboa, Portugal). A total of 500 ng of RNA was reverse-transcribed into cDNA using the Maxima First Strand cDNA Synthesis kit from Thermo Scientific (Thermo Fisher Scientific, Massachusetts, USA). Quantitative real-time PCR (RT-qPCR) analysis was carried out using a Roche LightCycler 480 II instrument (Roche Diagnostics, Barcelona, Spain).

#### Yeast Two-Hybrid (Y2H) assay

Pray and bait expression clones were produced using the Gateway Cloning System (Thermo Fisher Scientific). Full-length coding sequences (CDS) of *MdBRC1*, *MdFT1* and *MdFT2* genes cloned into pDONR201 were transferred by LR reaction (Thermo Fisher Scientific) into the destination vectors pDEST22 and pDEST32. All vectors were confirmed by sequencing. Yeast PJ69-4A strain ^66^ was used for co-transformation according to the Frozen-EZ Yeast Transformation II™ protocol. The different combinations of binding domain and activation domain plasmids were spread on SD plates lacking Leu and Trp (SD/-Leu/-Trp) medium plates, and incubated at 30°C for 3– 4 days for yeast selection. Positive clones were then inoculated into selective media lacking Leu, Trp and His (SD/-His/-Leu/-Trp) for interaction detection. The baits containing genes were tested for autoactivation in SD/-His/-Leu/-Trp plates supplemented with 0.8 mM 3-amino-1,2,4-triazole (3-AT), incubated at 30°C for approximately 4 days. All interactions were confirmed by sequencing.

## QUANTIFICATION AND STATISTICAL ANALYSIS

Statistical analyses were done with GraphPad Prism 7.0 software. The statistical details of experiments can be found in the corresponding Figure legends.

## Acknowledgements

Helena Gioppato was the recipient of a Campus France Fellowship (reference 148405U). This project received funding from ERA-NET SusCrop2 (FruitFlow, ANR-21-SUSC-0002). MGX acknowledges financial support from France Génomique National infrastructure, funded as part of the “Investissement d’Avenir” program managed by Agence Nationale pour la Recherche (contract ANR-10-INBS-09).

## Author contributions

Helena Augusto Gippato contributed to investigation, project administration and writing. Joan Estevan contributed to investigation and writing. Mohamad Al Bolbol contributed to investigation. Alexandre Soriano contributed to formal analysis. Julio Garighan contributed to investigation. Kwanho Jeong contributed to investigation. Céline Georget contributed to investigation. Daniela Gómez-Soto contributed to investigation and writing. Samer El Khoury contributed to formal analysis. Vítor da Silveira Falavigna contributed to investigation, Simon George contributed to formal analysis. Mariano Perales contributed to investigation, supervision and writing. Fernando Andrés contributed to conceptualisation, funding acquisition, project administration, supervision and writing.

