## Supplementary Figures and Legends for "MdBRC1 and MdFT2 Interaction Fine-Tunes Bud Break Regulation in Apple"

### SUPPLEMENTARY FIGURE LEGENDS

**Supplementary Figure 1.** Phylogenetic analysis of the TCP transcription factor family from *Arabis alpina* (Aa), *Arabidopsis thaliana* (At), *Malus domestica* (Md) and *Populus trichocarpa* (Pt). The unrooted tree was constructed based on the full-length protein sequences using the maximum likelihood method. TCP proteins are grouped into two major classes: Class I (orange) and Class II (green), with Class II further subdivided into the CIN (light blue) and CYC/TB1 (dark blue) subclades. The PFC (Proliferation Factor Clade), indicated in dark red, highlights a specific subgroup within Class I. Bootstrap support values, assessed using 1000 replicates, are shown at key nodes. The blue circle indicates *BRC1* orthologues.

**Supplementary Figure 2. (A)** Alignment of the two apple TCP18 paralogs and the AtBRC1 sequences; **(B)** Alignment of the two apple TCP18 paralogs; **(C)** Alignments and sequence similarity analysis between the MdTCP18a and AtBRC1; **(D)** Alignment and sequence similarity analysis between the MdTcp18b and AtBRC1. Pairwise sequence identity and similarity were calculated using the Sequence Manipulation Suite – Ident and Sim tool, based on aligned sequences in FASTA format.

**Supplementary Figure 3.** Characterization of MdBRC1 transgenic lines. Relative expression analysis of MdBRC1 lines by qRT-PCR. *Ubiquitin7* was used as the housekeeping gene. Plotted values and error bars are fold-change means ± s.d. of two biological replicates.

**Supplementary Figure 4.** Relative expression analysis of ABA and GA-related genes in MdBRC#8 and MdBRC#9 apical buds. Values represent the mean expression from two apical buds. Significant differences among lines and the WT were determined by one-way ANOVA followed by a Tukey test, $*p < 0.05$.

**Supplementary Figure 5.** Relative expression levels of MdTCP14a and MdTCP15b across dormancy at the beginning and end of endodormancy (C and O, respectively), measured by RNA-seq. Data are presented as mean TPM (Transcripts Per Million) ± SEM ($n = 3$ biological replicates).

**SUPPLEMENTARY FIGURES**

**Supplementary Figure 1**

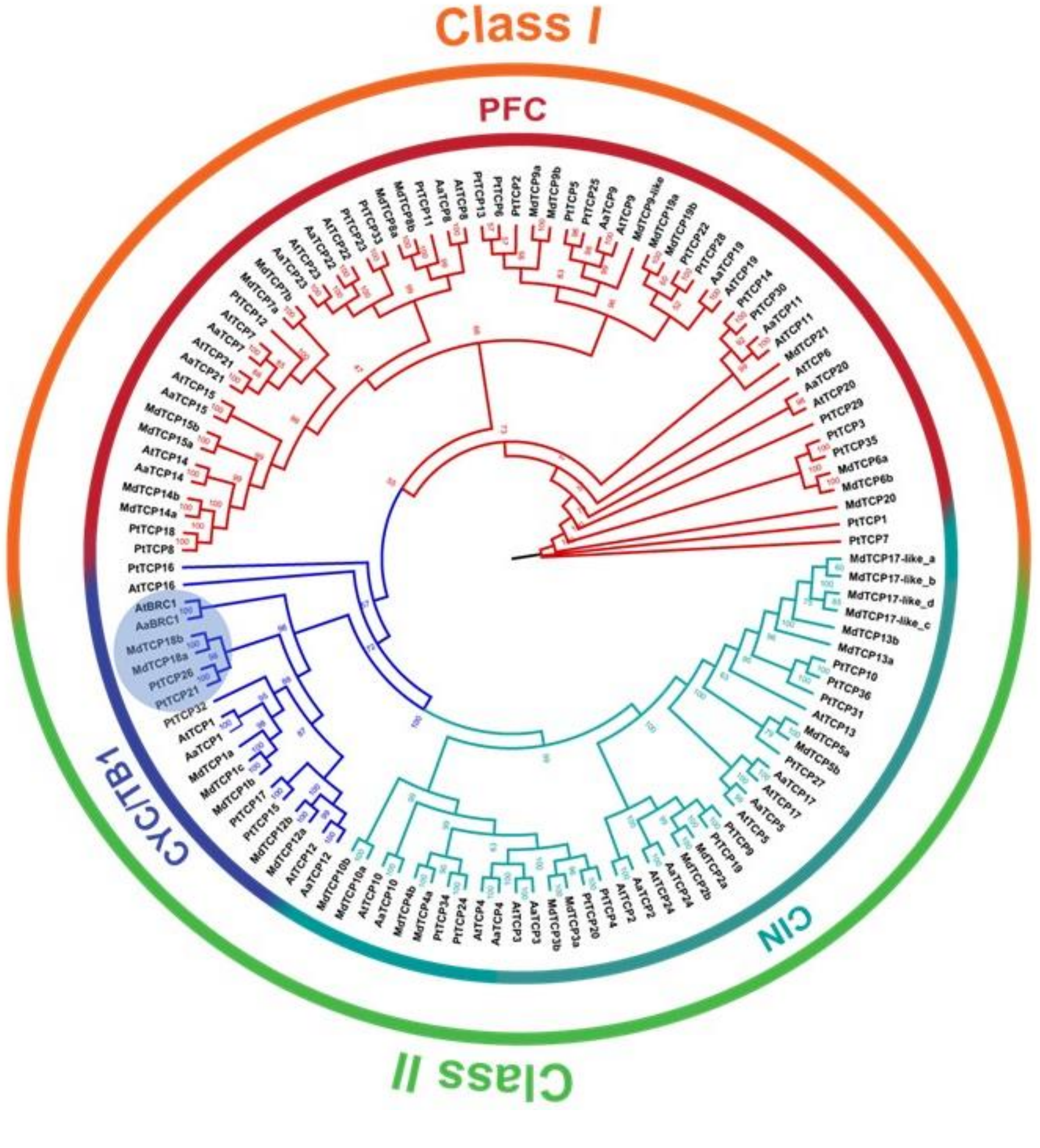

### Supplementary Figure 2A

Color Align Conservation results

```
MdTCP18a MFPSHN-NNLNELVPVSYPHVDQSIFHSWPSYHDNSTLTPNSLTILNPNPNS-RQQEGGEENLHQQH-HHLPFSLLYFPS 77
MdTCP18b MFPSDNTSNVNELIPVSYPHVDQPFFHSRPFDHDITTVTPNFFTNSNPNPNSGQQQEGGEENLHQQH-HHPPLSLLYFPS 79
AtBRC1   ---------------------------------------------MNNNIFSTTTTINDDYMLFPYNDHYSSQPLLPFSP 35

MdTCP18a PFEDDDVLLFDQQHHHQEPDHIELSIHESQEPLYFMKEAAAAA----------------AADDNT--VVGNDHQKTTTTS 139
MdTCP18b PFEDDDVLLF-QQHHHQEPDHIGISLHESQVPPYFMKEAAAAATATAAAAAAAAAAAATAADDNTGTVVGVDHQKTTTTS 158
AtBRC1   SSSINDILIHSTSNTSNNHLDHHHQFQQPSPFSHFEFAPDCALLTSFHPENNGHDDNQTIPNDNHHPSLHFPLNNTIVEQ 115

MdTCP18a VNRKMVDWDSNKKGHRMNMDDQPQIPR-KRTSKRDRHSKINTARGPRDRRMRLSLEVALKFFGLQDALGFDKPSKTVEWL 218
MdTCP18b VNIKMVDWDSNKNGHHVYMDDQPQIPR-RRTSKRDRHSKINTARGPRDRRMRLSLEVARKFFGLQDVLEFDKASKTVEWL 237
AtBRC1   PTEPSETINLIEDSQRISTSQDPKMKKAKKPSRTDRHSKIKTAKGTRDRRMRLSLDVAKELFGLQDMLGFDKASKTVEWL 195

MdTCP18a LIQSEPAIKKLSR--DHHRQFNY--KHMVRCAKSTSPATSESCEVLS--GVDEAPTNVNISING-----KVRSRGIKPSA 287
MdTCP18b LIQSEPEIKKLSR--DHHRKFNY--KNMVRCAKTTSPATSESCEMLS--GVDEAPTNINISNGGNDDNDKVRSSGIKPSA 311
AtBRC1   LTQAKPEIIKIATTLSHHGCFSSGDESHIRPVLGSMDTSSDLCELASMWTVDDRGSNTNTT--------ETRGNKVDGRS 267

MdTCP18a KERKFVHRQSRKSAFHPLAKASREKARARARERTREKMQRSKKPSNDQANSSRLSSWNPFETEEESPAHNNNMNSTTNDQ 367
MdTCP18b KERKIVHRQSRKGAFHPLAKASREKARARARERTREKMQRSKKPSNDQAKLSRLNSWNPFESGEESSAYNNNMNNTTNDQ 391
AtBRC1   MRGKRKRPEPRTPILKKLSKEERAKAREERAKGRTMEKMMMKMKGRSQLVKVVEEDAHDHGEIIKNNN--RSQVNRSSFEM 345

MdTCP18a PNSRAVLRDYPDEVQEPLSSPQAGNIQDMVVDHGTTHDAMVVLGKWSPPPVFTRLQQNTGISQEHQQFAD--FQFFGKPW 445
MdTCP18b PDSMVALRPYPDGVEEPLSSSLAGDIQDMVVDHGTTHDAMMVMGKWSPSSVFTPLQQNTGISQEHQQFAD--FQFFDKPW 469
AtBRC1   THCEDKIEELCKNDRFAVCNEFIMNKKDHISNE--SYDLVNYKPNSSFPVINHHRSQGAANSIEQHQETDLHYSFGAKPR 423

MdTCP18a EVDNNTHNLF 455
MdTCP18b DVYNNTHKLF 479
AtBRC1   DLMHNYQNMY 433
```

### Supplementary Figure 2B

Color Align Conservation results

```
MdTCP18a MFPSHN-NNLNELVPVSYPHVDQSIFHSWPSYHDNSTLTPNSLTILNPNPNS-RQQEGGEENLHQQHHHLPFSLLYFPSP 78
MdTCP18b MFPSDNTSNVNELIPVSYPHVDQPFFHSRPFDHDITTVTPNFFTNSNPNPNSGQQQEGGEENLHQQHHHPPLSLLYFPSP 80

MdTCP18a FEDDDVLLFDQQHHHQEPDHIELSIHESQEPLYFMKEAAAAA----------------AADDNT--VVGNDHQKTTTTSV 140
MdTCP18b FEDDDVLLF-QQHHHQEPDHIGISLHESQVPPYFMKEAAAAATATAAAAAAAAAAAATAADDNTGTVVGVDHQKTTTTSV 159

MdTCP18a NRKMVDWDSNKKGHRMNMDDQPQIPRKRTSKRDRHSKINTARGPRDRRMRLSLEVALKFFGLQDALGFDKPSKTVEWLLI 220
MdTCP18b NIKMVDWDSNKNGHHVYMDDQPQIPRRRTSKRDRHSKINTARGPRDRRMRLSLEVARKFFGLQDVLEFDKASKTVEWLLI 239

MdTCP18a QSEPAIKKLSRDHHRQFNYKHMVRCAKSTSPATSESCEVLSGVDEAPTNVNISING-----KVRSRGIKPSAKERKFVHR 295
MdTCP18b QSEPEIKKLSRDHHRKFNYKNMVRCAKTTSPATSESCEMLSGVDEAPTNINISNGGNDDNDKVRSSGIKPSAKERKIVHR 319

MdTCP18a QSRKSAFHPLAKASREKARARARERTREKMQRSKKPSNDQANSSRLSSWNPFETEEESPAHNNNMNSTTNDQPNSRAVLR 375
MdTCP18b QSRKGAFHPLAKASREKARARARERTREKMQRSKKPSNDQAKLSRLNSWNPFESGEESSAYNNNMNNTTNDQPDSMVALR 399

MdTCP18a DYPDEVQEPLSSPQAGNIQDMVVDHGTTHDAMVVLGKWSPPPVFTRLQQNTGISQEHQQFADFQFFGKPWEVDNNTHNLF 455
MdTCP18b PYPDGVEEPLSSSLAGDIQDMVVDHGTTHDAMMVMGKWSPSSVFTPLQQNTGISQEHQQFADFQFFDKPWDVYNNTHKLF 479
```

**Results for MdTCP18a vs MdTCP18b:**
Alignment length: 480
Identical residues: 381
Similar residues: 18
Percent identity: 79.38
Percent similarity: 83.13

### Supplementary Figure 2C

Color Align Conservation results

```
MdTCP18a MFPSHNNNLNELVPVSYPHVDQSIFHSWPSYHDNSTLTPNSLTILNPNPNSRQQEGGEENLHQQHHHLPFSLLYFPSPFE 80
AtBRC1   -------MNNNIFSTTTTINDDYMLFPYNDHYSSQPLLPFSPSS-SINDILIHSTSNTSNNHLDHHHQ----FQQPSPFS 68

MdTCP18a DDDVLLFDQQHHHQEPDHIELSIHESQEPLYFMKEAAAAAAADDNTVVGNDHQKTTTTSVNRKMVDWDSNKKGHRMNMDD 160
AtBRC1   HFE---FAPDCALLTSFHPENNGHDDNQTIPNDNHHPSLHFPLNNTIVEQPTEPSETINLI---------EDSQRISTSQ 136

MdTCP18a QPQIPR-KRTSKRDRHSKINTARGPRDRRMRLSLEVALKFFGLQDALGFDKPSKTVEWLLIQSEPAIKKLSR--DHHRQF 237
AtBRC1   DPKMKKAKKPSRTDRHSKIKTAKGTRDRRMRLSLDVAKELFGLQDMLGFDKASKTVEWLLTQAKPEIIKIATTLSHHGCF 216

MdTCP18a NY--KHMVRCAKSTSPATSESCEVLS--GVDEAPTNVNISINGKVRSRGIKPSAKERKFVHRQSRKSAFHPLAKASREKA 313
AtBRC1   SSGDESHIRPVLGSMDTSSDLCELASMWTVDDRGSNTNTT---ETRGNKVDGRSMRGKRKRPEPRTPILKKLSKEERAKA 293

MdTCP18a RARARERTREKMQRSKKPSNDQANSSRLSSWNPFETEEESPAHNNNMNSTTNDQPNSRAVLRDYPDEVQEPLSSPQAGNI 393
AtBRC1   RERAKGRTMEKMMMKMKGRSQLVKVVEEDAHDHGEIIKNN--NRSQVNRSSFEMTHCEDKIEELCKNDRFAVCNEFIMNK 371

MdTCP18a QDMVVDHGTTHDAMVVLGKWSPPPVFTRLQQNTGISQEHQQFAD--FQFFGKPWEVDNNTHNLF 455
AtBRC1   KDHISNE--SYDLVNYKPNSSFPVINHHRSQGAANSIEQHQFTDLHYSFGAKPRDLMHNYQNMY 433
```

**Results for MdTCP18a vs AtBRC1:**
Alignment length: 464
Identical residues: 116
Similar residues: 70
**Percent identity: 25.00**
**Percent similarity: 40.09**

### Supplementary Figure 2D

Color Align Conservation results

```
AtBRC1   ---------------------------------------------MNNNIFSTTTTINDDYMLFPYNDHYSSQPLLPFSP 35
MdTCP18b MFPSDNTSNVNELIPVSYPHVDQPFFHSRPFDHDITTVTPNFFTNSNPNPNSGQQQEGGEENLHQQHHHPPLSLLYFPSP 80

AtBRC1   SSSINDILIHSTSNTSNNHLDHHHQFQQPSPFSHFEFAPDCALLTSFHPENNGHDDNQTIPNDNHHPSLHFPLNNTIVEQ 115
MdTCP18b FEDDDVLLFQQHHHQEPDHIGISLHESQVPPYFMKEAA--AAATATAAAAAAAAAAATAADDNTGTVVGVDHQKTTTTS 158

AtBRC1   PTEPSETINLIEDSQRISTSQDPKMKKAKKPSRTDRHSKIKTAKGTRDRRMRLSLDVAKELFGLQDMLGFDKASKTVEWL 195
MdTCP18b VNIKMVDWDSNKNGHHVYMDDQPQIPR-RRTSKRDRHSKINTARGPRDRRMRLSLEVARKFFGLQDVLEFDKASKTVEWL 237

AtBRC1   LTQAKPEIIKIATTLSHHGCFSSGDESHIRPVLGSMDTSSDLCELASMWTVD------DRGSNTNTTETRGNKVDGRSMR 269
MdTCP18b LIQSEPEIKKLSR--DHHRKFNY--KNMVRCAKTTSPATSESCEMLSGVDEAPTNINISNGGNDDNDKVRSSGIKPSAKE 313

AtBRC1   GKRKRPEPRTPILKKLSKEERAKARERAKGRTMEKMMMKMKGRSQLVKVVEEDAHDHGEIIKNNN--RSQVNRSSFEMTH 347
MdTCP18b RKIVHRQSRKGAFHPLAKASREKARARARERTREKMQRSKKPSNDQAKLSRLNSWNPFESGEESSAYNNNMNNTTNDQPD 393

AtBRC1   CEDKIEELCKNDRFAVCNEFIMNKKDHISNE--SYDLVNYKPNSSFPVINHHRSQGAANSIEQHQFTDLHYSFGAKPRDL 425
MdTCP18b SMVALRPYPDGVEEPLSSSLAGDIQDMVVDHGTTHDAMMVMGKWSPSSVFTPLQQNTGISQEHQQFAD--FQFFDKEWDV 471

AtBRC1   MHNYQNMY 433
MdTCP18b YNNTHKLF 479
```

**Results for AtBRC1 vs MdTCP18b:**
Alignment length: 488
Identical residues: 108
Similar residues: 64
**Percent identity: 22.13**
**Percent similarity: 35.25**

**Supplementary Figure 3**

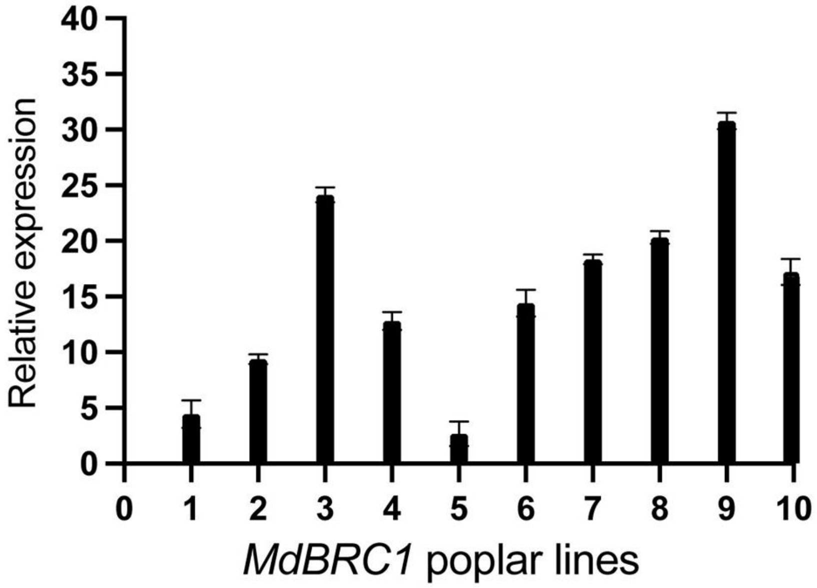


**Supplementary Figure 4**

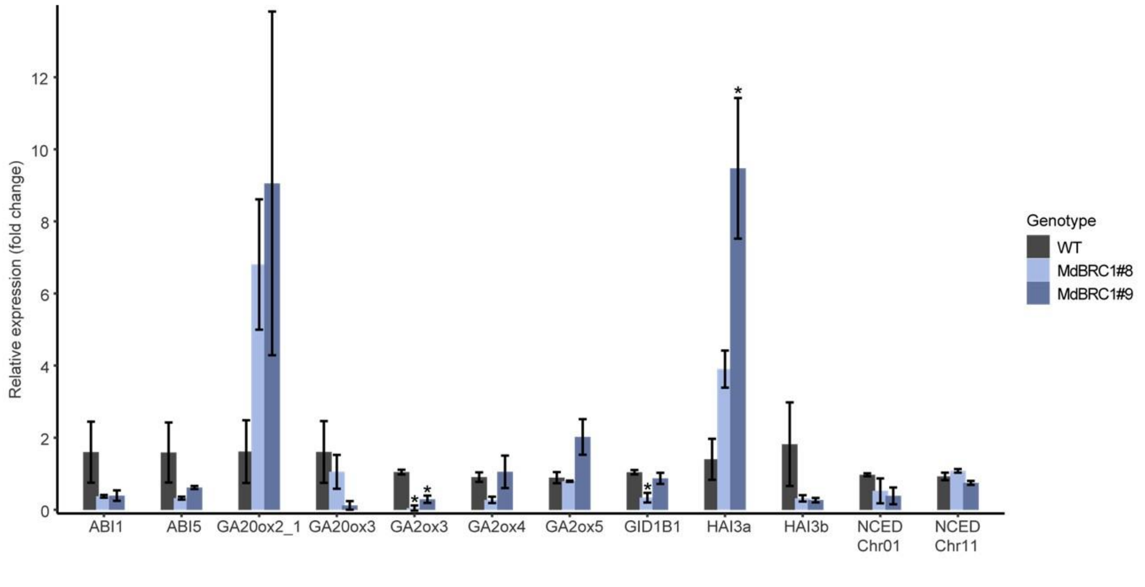

**Supplementary Figure 5**

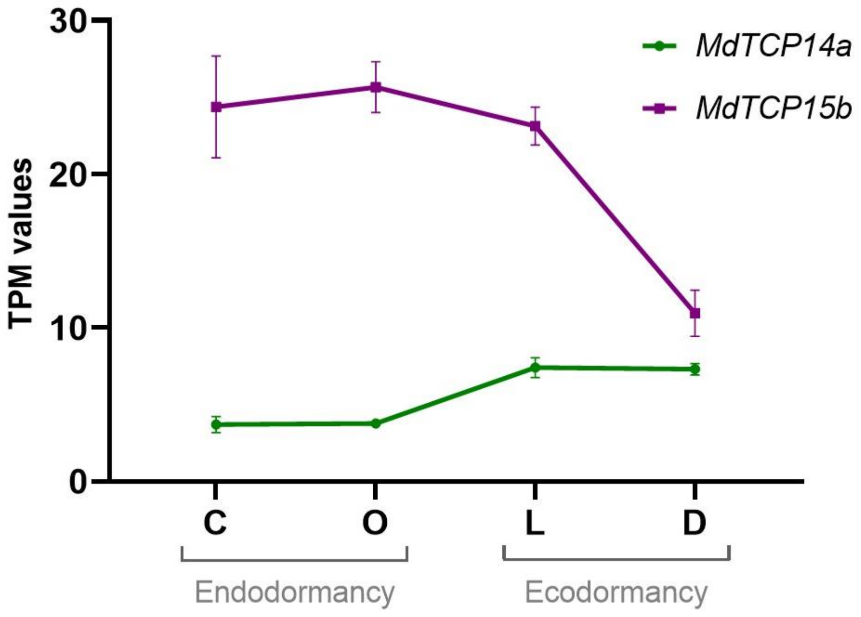
